# Cancer Cell Line Profiler (CCLP): a webserver for the prediction of compound activity across the NCI60 panel

**DOI:** 10.1101/105478

**Authors:** Isidro Cortés-Ciriano, Daniel S. Murrell, Bernard Chetrit, Andreas Bender, Thérèse Malliavin, Pedro J. Ballester

## Abstract

**Summary:** CCLP (**C**ancer **C**ell **L**ine **P**rofiler) is a webserver for the prediction of compound activity across the NCI60 panel. CCLP uses a multi-task Random Forest model trained on 941,831 data-points that integrates structural information from 17,142 compounds and multi-omics data sets from 59 cancer cell lines. In addition, CCLP also implements conformal prediction to provide individual prediction errors at several confidence levels. CCLP computes compound descriptors for a set of input molecules and predicts their activity across the NCI60 panel. The output of running CCLP consists of one barplot *per* input compound displaying the predicted activities and errors across the NCI60 panel, as well as a text file reporting the predicted activities and errors in prediction

**Availability:** CCLP is freely available on the web at cclp.marseille.inserm.fr

## 1 Introduction

Although cultured cancer cell lines do not fully recapitulate the complex tumor microenvironment, they have proved versatile preclinical models to study the pharmacology of anticancer drugs (Iorio, 2016). Multiple initiatives have characterized thousand of cell lines at the molecular level, as well as their response to large collections of small molecules (Barretina, 2012; Iorio, 2016; de Waal, 2016). The Developmental Therapeutics Program (DTP) from the United States National Cancer Institute (NCI) pioneered these efforts in the early 1990s by assembling a collection of 59 cancer cell lines spanning 9 cancer types (Shoemaker, 2006). This increasing wealth of in vitro sensitivity data has fostered the development of drug sensitivity prediction algorithms. The ultimate aim of these models is to predict the best pharmacological treatment on the basis of the patients’ genomic makeup. Although much remains to be done to realise this vision, the computational integration of pharmacological and genomic data of cancer cell lines has proved useful to investigate the effect of genomic variability on compound activity, and to model the *in vitro* activity of anticancer drugs (Cortés-Ciriano, 2016). Building machine learning models to predict compound activity generally requires expert curation and preparation of large amounts of data. While a plethora of studies have reported predictive models of compound activity on cell lines using both single-and multi-task learning strategies, reviewed in (Cortés-Ciriano *et al.*, 2016), no tool has been developed to date to enable the broader scientific community to use drug sensitivity prediction algorithms trained on the NCI60 data without requiring user knowledge of machine learning or descriptor generation. To fill this gap we have developed an *ad hoc* webserver, namely the Cancer Cell Line Profiler (CCLP). CCLP simplifies the use of drug sensivity prediction algorithms by eliminating the need for downloading and processing cell line sensitivity data, as it only requires a set of molecules as input, whose activity across the NCI60 panel is predicted in the back end. To facilitate the visualization of the predictions, CCLP also reports the results as a barplot accompanied by individual errors in prediction calculated with conformal prediction.

## 2 Results and performance

### 2.1 Cancer cell line sensitivity prediction

The models hosted by CCLP were trained on a data set comprising 941,831 50% growth inhibition bioassay end-points (GI50) of 17,142 compounds screened against the NCI60 panel (matrix 93.08% complete) (Cortés-Ciriano, 2016). We have previously shown that the integration of chemical and biological information from multiple compounds and cell lines in multi-task learning models improves the prediction of compound activity with respect to single-task models, where the activity of single compounds is modelled independently. Therefore, we decided to use the multi-task learning paradigm *proteochemometrics* to train the core models hosted in the CCLP (Menden *et al.*, 2013; Cortés-Ciriano *et al.*, 2015). Compounds were encoded with Morgan fingerprints given their high performance in drug sensitivity prediction studies, whereas cell lines were encoded with the average expression of the 1,000 canonical pathways displaying the highest variability across the NCI60 panel. Each compound-cell line pair was encoded by the concatenation of these two types of descriptors. This modelling framework, resulting from throrough optimization of model parameters and feature selection, has been shown to provide robust performance in cross-validation, with RMSE values in the 0.56-0.58 pGI50 units range (Cortés-Ciriano, 2016). Further assessment of this model revealed high power to extrapolate the training data to new cell lines and, although to a lesser extent, to new compounds as well, whereas the drug-pathway associations identified with the experimental data could be mostly retrieved using the cross-validation predictions.

### 2.2 Calculation of confidence intervals with conformal prediction

In conformal prediction (Norinder, 2014), the similarity (*i.e.* conformity) of a new data point to those used for training is quantified with a nonconformity score, *e.g*. 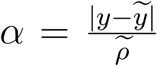, where *y* and 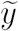 are the observed and the predicted values, respectively, calculated with a point prediction RF model, and 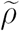 is the predicted error calculated with *e.g.* an RF error model. The point prediction model was generated using 10-fold cross validation on compound and cell-line descriptors as covariates, and pGI50 values as the dependent variable. The cross validation residuals of this model 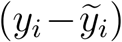 served as the depedent variable to train the error model, which was trained on the same covariates as the point prediction model. The cross-validation predictions from these two models were used to generate the vector of nonconformity scores for the training set, which after being sorted in increasing order is defined as: 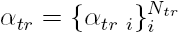 where *N*_*tr*_ is the number of datapoints in the training set. The value associated to the user-defined confidence level, *α*_*∈*_, is calculated as: *α*_*∈*_ = *α*_*tr i*_ *if i* ≡ |*N*_*tr*_ * *∈*| where ≡ indicates equality. Next, the errors in prediction, 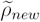, and the predicted pGI50 values, 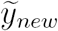, for a new data point (*x*_*new*_) are predicted with the error and the point prediction models, respectively. The individual confidence interval (CI) for *x*_*new*_ is defined as 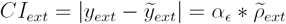. The confidence region (CR) is finally defined as: 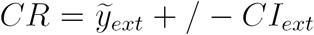. The interpretation of the confidence regions is straightforward. For instance, a confidence level of 80% means that the true pGI50 value will lie outside the predicted confidence region in at most 20% of the cases.

## 3 Implementation

The CCLP site has been implemented using the Django web framework (https://djangoproject.com), whereas the core CCLP models have been developed using the Python programming language and the scikit learn library (Pedregosa, 2011). To run CCLP the users need to upload a single molecule file in smiles or sdf format. For those input molecules passing a validity check, CCLP computes Morgan fingerprints, which are further assembled with precomputed cell line descriptors. These composite descriptors are input to the core CCLP models to These predictions, as well as a barplot representation thereof, are emailed to the user.

## 4 Conclusion

We present a webserver for the prediction of compound activity on the NCI60 panel. Given that the predictive capabilities of machine learning models are restricted by the training data, we provide individual errors in prediction that quantify the reliability of each individual prediction and are easy to interprete. CCLP is designed to be useful to the broader scientific community, as users with no experience in machine learning or bioactivity data curation can easily obtain predictions for small molecules by uploading a single molecule file. CCLP runs in a reasonable amount of time (~10 seconds and ~8 minutes for 1 and 1,000 molecules, respectively), and the performance of its core models has been shown to be competitive with the state of the art. In the future, we plan to make available additional drug sensitivity prediction models trained on other data sets and extend the models currently hosted by the CCLP as more cell line sensitivity and multi-omics data become publicly available.

## Acknowledgements

The authors acknowledge the computational resources provided by the Institut Pasteur and the Centre de Recherche en Cancérologie de Marseille (CRCM). ICC and TM thank Institut Pasteur and CNRS for funding. This project has received funding from the European Union’s Framework Programme For Research and Innovation Horizon 2020 (2014-2020) under the Marie Curie Skłodowska-Curie Grant Agreement No. 703543. AB thanks the European Research Commission Starting Grant ERC-2013-StG 336159 MIXTURE.

